# Insufficient fumarase contributes to generating reactive oxygen species in Dahl salt sensitive rats

**DOI:** 10.1101/302182

**Authors:** Entai Hou, Xuewei Zheng, Zhe Yang, Xian Li, Zerong Liu, Meng Chen, Xiaoxue Li, Mingyu Liang, Zhongmin Tian

**Affiliations:** The Key Laboratory of Biomedical Information Engineering of Ministry of Education, School of Life Science and Technology, Xi’an Jiaotong University, Xi’an 710049, China; Center of Systems Molecular Medicine, Department of Physiology, Medical College of Wisconsin, Milwaukee, WI 53226

**Author notes:** These authors contributed equally to this work (Entai Hou, Xuewei Zheng). Corresponding author: Dr. Zhongmin Tian, School of Life Science and Technology, Xi’an Jiaotong University, Xi’an 710049, China.

**Keywords:** Dahl salt sensitive rats, Fumarase, Fumarate, Reactive oxygen species, Glutathione metabolism

## Abstract

Dahl SS rats exhibit greater levels of renal medullary oxidative stress and lower levels of fumarase activities. Fumarase insufficiencies can increase reactive oxygen species (ROS), the mechanism of which, however, is not clear. A proteomic analysis indicated fumarase knockdown in HK-2 cells resulted in changes in the expression or activity of NADPH oxidase, mitochondrial respiratory chain complex I and III, ATP synthase subunits, and α-oxoglutarate dehydrogenase, all of which are sites of ROS formation. Meantime, the activities of key antioxidant enzymes such as G6PD, 6PGD, GR, GPx and GST increased significantly too. The apparent activation of antioxidant defense appeared insufficient as glutathione precursors, glutathione and GSH/GSSG ratio were decreased. SS rats exhibited changes in redox metabolism similar to HK-2 cells with fumarase knockdown. Supplementation with fumarate and malate, the substrate and product of fumarase, increased and decreased, respectively, blood pressure and the levels of H_2_O_2_ and MDA in kidney tissues of SS rats. These results indicate fumarase insufficiencies cause a wide range of changes at several sites of ROS production and antioxidant mechanisms.

## Introduction

Salt-sensitive hypertension, a multifactorial disease affecting nearly 15% of the world population, is a major risk factor for stroke, heart failure, and end-stage renal disease [1-3]. Dahl salt sensitive (SS) rat, as an animal model of human salt sensitive hypertension, was widely used to dissect the pathogenesis. SS.13^BN^ rats as a normotensive control, which chromosome 13 of the Brown Norway (BN) rat was integrated into the genetic background of the SS and result in a significant reduction of blood pressure and renal injury [4], had tremendously narrowed the interested gene on chromosome 13. There is only a 1.95% allelic difference over the entire genome between SS and SS.13^BN^ rats. Meanwhile, SS rats renal medullary exhibited obviously oxidative stress and blood pressure salt-sensitivity compared with SS.13^BN^ rats [5], as H_2_O_2_ concentrations had also been found to be nearly twice as great in the dialysate of SS rats when both rats were maintained on a 0.4% NaCl diet [6]. The level was more than doubled with a 4% NaCl in diet, but it remained significantly higher in SS rats [6]. Furthermore, the increased production of O_2_.^-^ in mTAL also could diffuses to surrounding vasa recta, contribute to decrease bioavailability of NO and reduce medullary blood flow, resulting in sustained hypertension in SS rats [7]. However, the mechanisms that excessive O_2_.^-^ and H_2_O_2_ produced in renal medulla of SS rats, is yet not fully understood.

Our previous differential proteomics study found that fumarase (FH), the gene of which locates on rat chromosome 13, was expressed differentially between SS and SS.13^BN^ rats, and the activity of FH is significantly lower in SS rats [8, 9]. As FH catalyzes the conversion between fumarate and L-malate, the insufficiency of FH activity in SS rat kidney was associated with increased levels of fumarate [8]. The intravenous infusion of a fumarate precursor diethyl-fumarate in SS.13^BN^ rats resulted in increased levels of fumarate in renal medullary and significantly exacerbated salt-induced hypertension in SS.13^BN^ rats [8]. Furthermore, excessive fumarate could also increase the level of H_2_O_2_ in vivo and in cultured HK-2 cells [8]. Our recent research work further demonstrated that overexpressing fumarase on the background of the SS rat, SS-TgFh1 transgenic rats, could significantly attenuate hypertension and H_2_O_2_ production [10]. So the present study will further dissect the imbalance of redox metabolism and understand the mechanism of ROS production by a proteomic analysis in fumarase knockdown HK-2 cells.

## Results

### Differentially expressed proteins between FH insufficient HK-2 cells and NC cells

The fumarase was knocked down by siRNA in HK-2 cell line, the protein profile of the whole cells was analyzed by iTRAQ based proteomics technology, and metabolites were detected by GC/MS. Content and activity of FH were examined to ensure the efficiency of FH knockdown. The sample preparation used for proteomic analysis same as the description we reported [11] and the sample test was showed in Figure EV1.

A total of 3174 distinct proteins were identified and quantified reliably at the value of global FDR less than 1% and identified with more than two peptides for 95% confidence. Details of the quantified proteins are shown in Table EV1. Compared with the NC, 267 identified proteins exhibited significant difference between FH knockdown HK-2 cells and NC cells (fold change >1.5, P<0.05) (Table EV2). Among them, 165 proteins displayed increased expression and 102 proteins displayed decreased expression compared to NC. SOD2, IDH1, IDH2 and HSP90 among the 267 differentially expressed proteins were further confirmed by Western blotting. As shown in Figure EV2, the relative protein content from western blotting was consistent with proteomic quantification.

### GO and KEGG analysis of the differential expressed proteins

To clarify the changed metabolic pathways that involved in the response to FH knockdown, 267 differential proteins were further examined using the Omics Bean bioinformatics tool. Gene ontology (GO) enrichment analysis was conducted in biological process (BP), cellular component (CC) and molecular function (MF) categories, respectively. An overview of significantly changed proteins in BP, CC and MF categories was shown in Figure EV3 and Table EV3. KEGG pathway-based analysis showed that ribosome, carbon metabolism, glutathione metabolism, mRNA transport, pyruvate metabolism, fatty acid degradation, glycolysis /gluconeogenesis and TCA cycle were significantly enriched among 267 differential proteins (Figure EV4 and Table EV4).

### The formation sites of excessive amounts of ROS and H_2_O_2_ in FH knockdown HK-2 cells

The activity of NOX were detected and showed significantly increased in FH insufficient HK-2 cells compared with NC cells (Figure 1A). The differential proteomic analysis data showed that expression of NADH dehydrogenase (ubiquinone) iron-sulfur protein 5 (NDUFS5), subunit of respiratory chain complex I, reduced for 1.5 times, while the expression of succinate dehydrogenase (Complex II) has no significant difference between FH knockdown HK-2 cells and NC cells. Rieske iron-sulfur protein (RISP) and Cyt b-c1 complex subunit 6 (Complex III) reduced 1.5 fold in FH knockdown HK-2 cells. Meanwhile, the expression of OGDH and B5R (cytochrome b5 reductase) increased 1.5 and 1.7 fold. Except these defined ROS generation sites, F_1_F_0_ ATP synthase subunits ATP5A1, ATP5C1, ATP5I and ATP5L reduced by 1.7, 1.4, 1.5 and 2.1 times and ATP level reduced significantly (Figure 1C). Briefly, NOX, α-OGDH, B5R, ATPase and Complex I and III would be the main sites of ROS formation in FH knockdown HK-2 cells.

**Figure 1.**
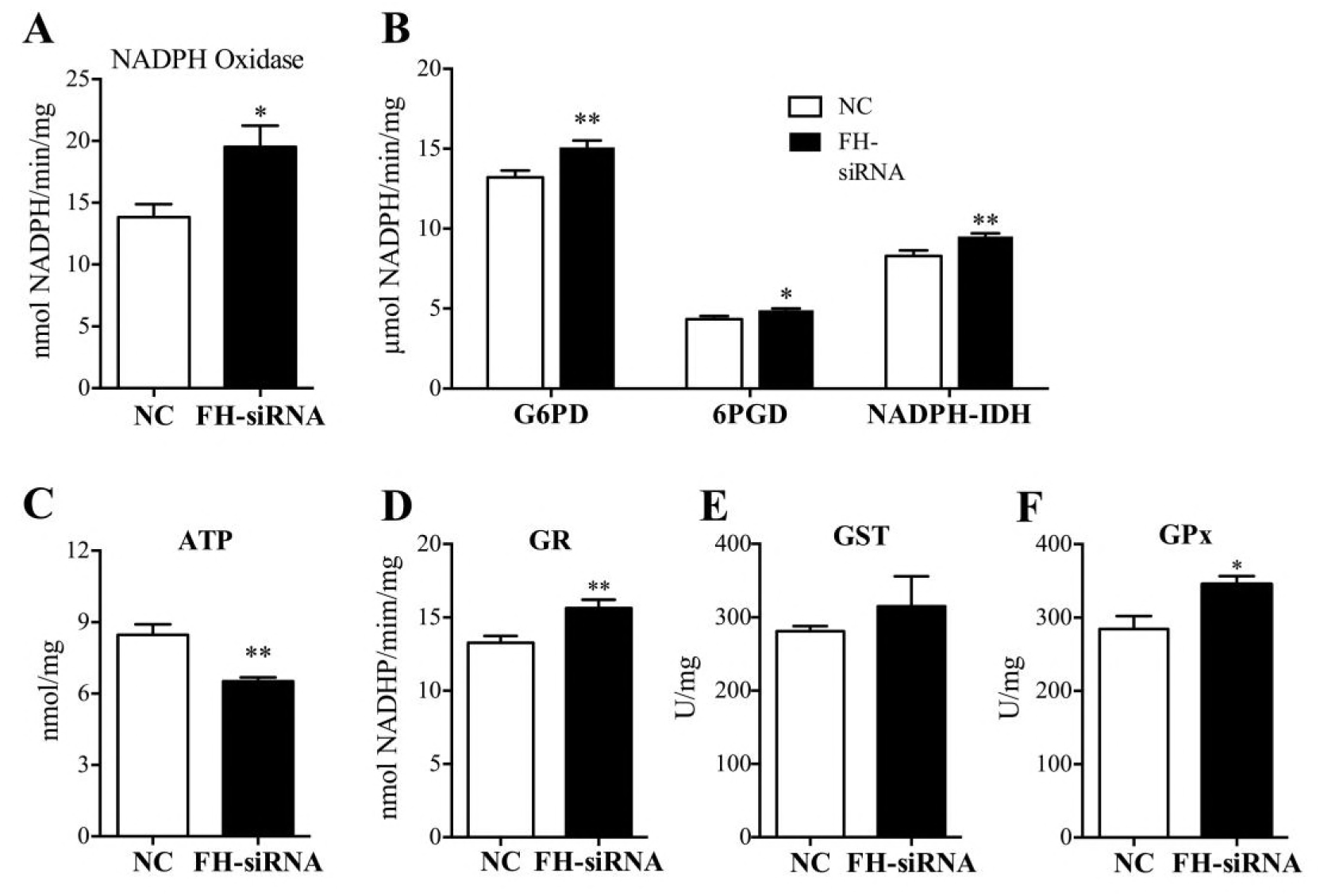
NADPH oxidase and key enzymes involved in glutathione metabolism activities in fumarase insufficient HK-2 cells. A NADPH oxidase activity enhanced in fumarase insufficiency HK-2 cells. B Fumarase insufficiency attenuated ATP production. C Glucose-6-phosphate 1-dehydrogenase (G6PD), 6-Phosphogluconate dehydrogenase (6PGD) and NADP^+^-IDH activity. D Glutathione reductase (GR) activity. E Glutathione S-transferase (GST) activity. F Glutathione peroxidase (GPx) activity. Mock siRNA targeting fumarase that had no sequence homology with any mammalian genes as a negative control. Data are presented as mean values ± SEM, n = 4, ^∗^p < 0.05, ^∗∗^p < 0.01 compared to negative control.

### Glutathione metabolism and antioxidant systems in FH knockdown HK-2 cells

The redox state of the cell is finely regulated by antioxidant systems, which include antioxidant enzymes and small molecular weight antioxidants. The proteomic analysis had identified the expression of all key enzymes involved in glutathione metabolism, increased their protein expression, which included Gpx, GR, GST, GS, G6PD and 6PGD (Table EV5), meanwhile, the activities of G6PD, 6PGD, GR, GST, GPx also increased compared with NC cells (Figure 1B, D, E, F). By contrast, there was no significant difference in the expression of glutamate cysteine ligase (GCL). In addition, the expression of other antioxidant enzymes, SOD2, NNTH (nicotinamide nucleotide transhydrogenase) and IDH2 increased 1.69, 1.77, 1.43 fold, however, the CAT, SOD1 had no significant difference between FH knockdown HK-2 cells and NC cells. The expression of SOD2, IDH1 and IDH2 was further confirmed by western blotting (Figure EV2). Furthermore, the expression of L-lactate dehydrogenase (LDHB), NADPH-dependent alcohol dehydrogenase (AKR1A1), glutathione dependent formaldehyde dehydrogenase (ADH5) and L-glutamate gamma semialdehyde dehydrogenase (ALDH7A1) increased 1.7, 1.5, 1.5, 1.4 fold compared with NC cells. These data suggested the increased activity of dehydrogenase would supply more antioxidants such as GSH, NADH or NADPH. Unfortunately, the content of GSH reduced dramatically, especially the ratio of GSH/GSSG decreased even if the content of NADPH increased compared with NC cells. The GSH synthesis substrate, glutamate, glycine and L-cysteine reduced significantly too (Figure 2 A-F).

**Figure 2.**
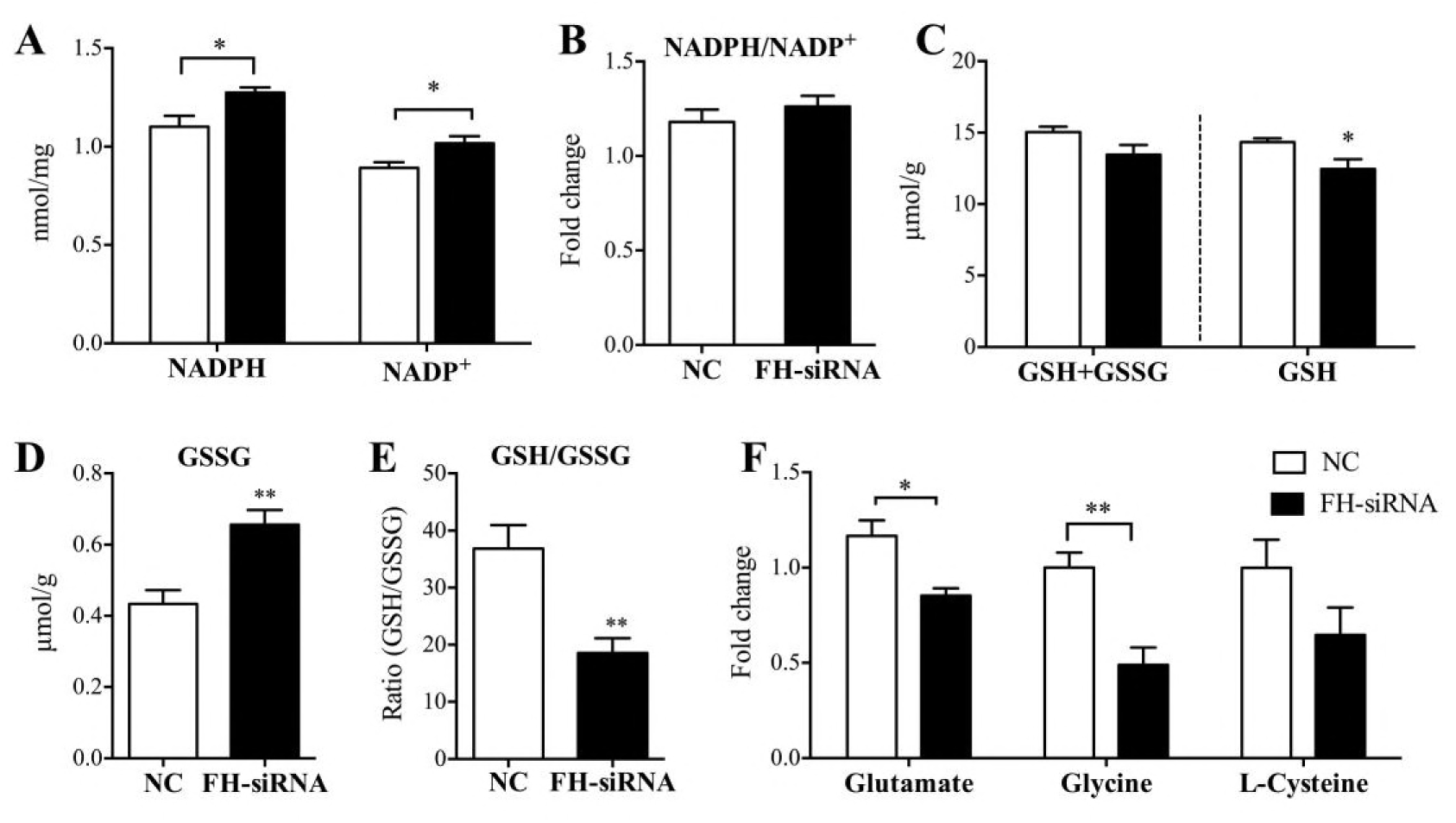
Fumarase insufficiency is associated with increased NADPH, NADP+ and reduced total GSH, GSH/GSSG ratio. A NADPH and NADP^+^ levels were detected in negative control and fumarase insufficient HK-2 cells. B NADPH/NADP^+^ ratio. C Total GSH (GSH+GSSH) and GSH levels. D GSSG levels. E GSH/GSSG ratio. F Fold changes of the glutamate, glycine and L-cysteine levels in fumarase insufficient HK-2 cells was calculated as a ratio relative to negative control. Mock siRNA targeting fumarase that had no sequence homology with any mammalian genes as a negative control. Data are presented as mean values ± SEM, n = 6, ^∗^p < 0.05, ^∗∗^p < 0.01 compared to negative control.

### Key enzymes involved in renal redox metabolism of Dahl SS rats

The FH activity was insufficient in renal cortex and medulla of SS rats compared with SS.13^BN^. To confirm that the renal cells exhibit the same response to the insufficient FH activity as FH knockdown HK-2 cells, the expression and activity of these key enzymes involved in redox metabolism were detected. For ROS formation, the activity of NOX had been reported increase definitely in renal medulla of SS rats [5], similar to FH knockdown HK-2 cells. The expression of OGDH increased significantly, and the expression of ATP5A1 reduced significantly in renal medulla (Figure 3). SS rats mitochondrial respiratory chain complex I and III had been reported involved in ROS production owing to the insufficient expression of EFTu (elongation factor Tu) protein in the mTAL mitochondria [12].

**Figure 3.**
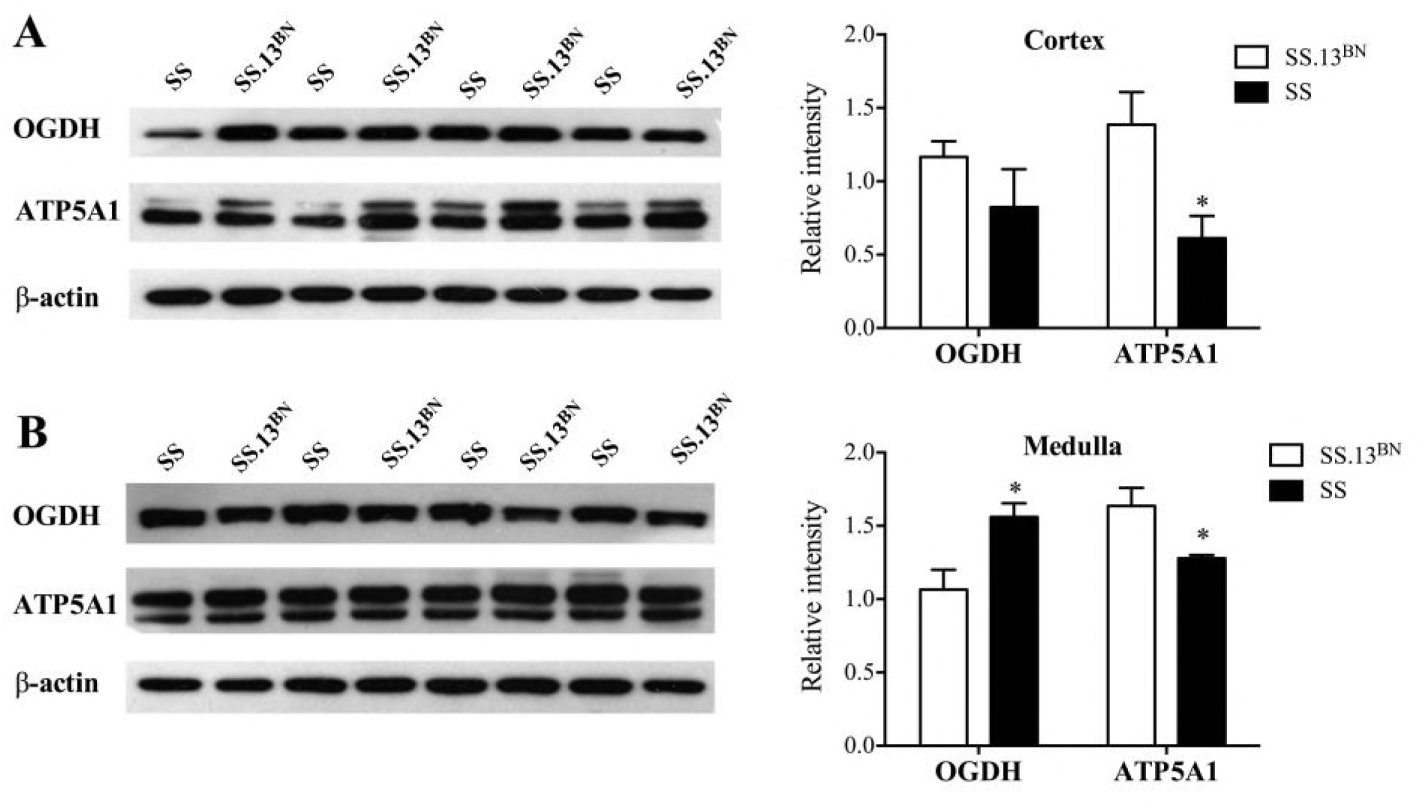
Western blot analysis of protein abundance of OGDH and ATP5A1 in the kidney of SS and SS.13^BN^ rats. A Abundance of α-oxoglutarate dehydrogenase (OGDH) and ATP synthase subunit alpha (ATP5A1) in the renal cortex. B Abundance of OGDH and ATP5A1 in the renal medulla. β-actin is used as the internal loading control. Data are presented as mean values ± SEM, n = 4, ^∗^p < 0.05 compared to negative control.

For the antioxidant response, the total GSH (GSH+GSSG), GSH reduced significantly in renal medulla (Figure 4A). The GSSG increased both in renal cortex and medulla (Figure 4B). The ratio of GSH/GSSG reduced significantly in renal medulla (P<0.01), and renal cortex (P=0.065) (Figure 4C). The GSH synthesis substrate glutamate, glycine and L-cysteine reduced significantly in renal medulla (Figure 4D). Meanwhile, the activity of G6PD increased in renal cortex and medulla, but the activity of 6PGD and NADP+-IDH only increased in renal cortex of SS rats compared with SS.^13^BN (Figure 4 E, F). These data suggested SS rats renal tissue had the similar redox metabolism to Fh knockdown HK2 cells.

**Figure 4.**
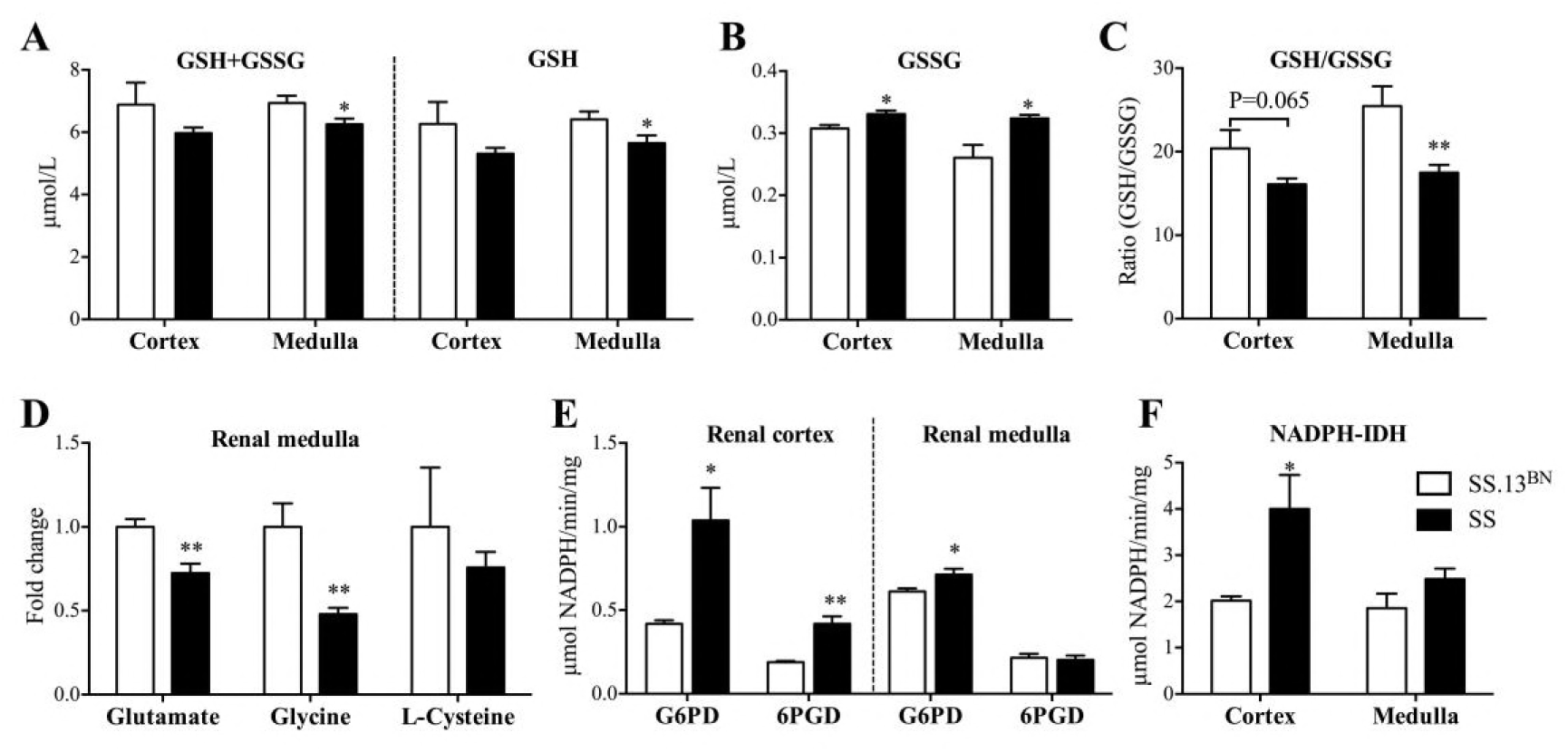
Total GSH levels and GSH/GSSG ratio were reduced in the kidney of SS rats. A Total GSH (GSH+GSSH) and GSH levels reduced in the renal cortex and medulla. B GSSG levels. C GSH/GSSG ratio. D Fold changes of the glutamate, glycine and L-cysteine in the renal medulla. E G6PD, 6PGD activitives in the renal cortex and medulla. F NADP^+^-IDH activity in the renal cortex and medulla. Data are presented as mean values ± SEM, n = 5, ^∗^p < 0.05 compared to negative control.

### Redox state in kidney with SS rat oral administration of fumarate and malate

The insufficiency of FH activity will result in the accumulation of fumarate and slight insufficiency of malate. As FH don’t directly involve in the formation of ROS, the metabolic intermediate fumarate or malate probably participate in the ROS production. To further verify the contribution of fumarate and malate to ROS generation, malate and fumarate were delivered by gavage in SS rats with normal salt diet and high salt diet. The level of H_2_O_2_ and MDA were detected after malate and fumarate supplementation for two weeks. The results showed the level of H_2_O_2_ and MDA increased significantly in renal cortex and medulla of SS rats with fumarate oral administration, while the blood pressure elevated significantly (Figure 5 A). On the contrary, oral administration of malate didn’t influence the level of H_2_O_2_ and MDA in SS rat with normal salt diet supplied and the blood pressure reduced significantly (Figure 5 B C). When SS rats fed with a high salt diet for two weeks, the level of H_2_O_2_ and MDA increased dramatically in renal medulla and the oral administration of malate could attenuate significantly the increasing of H_2_O_2_, MDA and blood pressure caused by high salt diet (Figure 5 D E). These data demonstrated that the substrate of FH, fumarate, could exacerbate ROS production and hypertension in SS rats, on the contrary, the product of FH, L-malate, could attenuate the hypertension induced by high salt diet.

**Figure 5.**
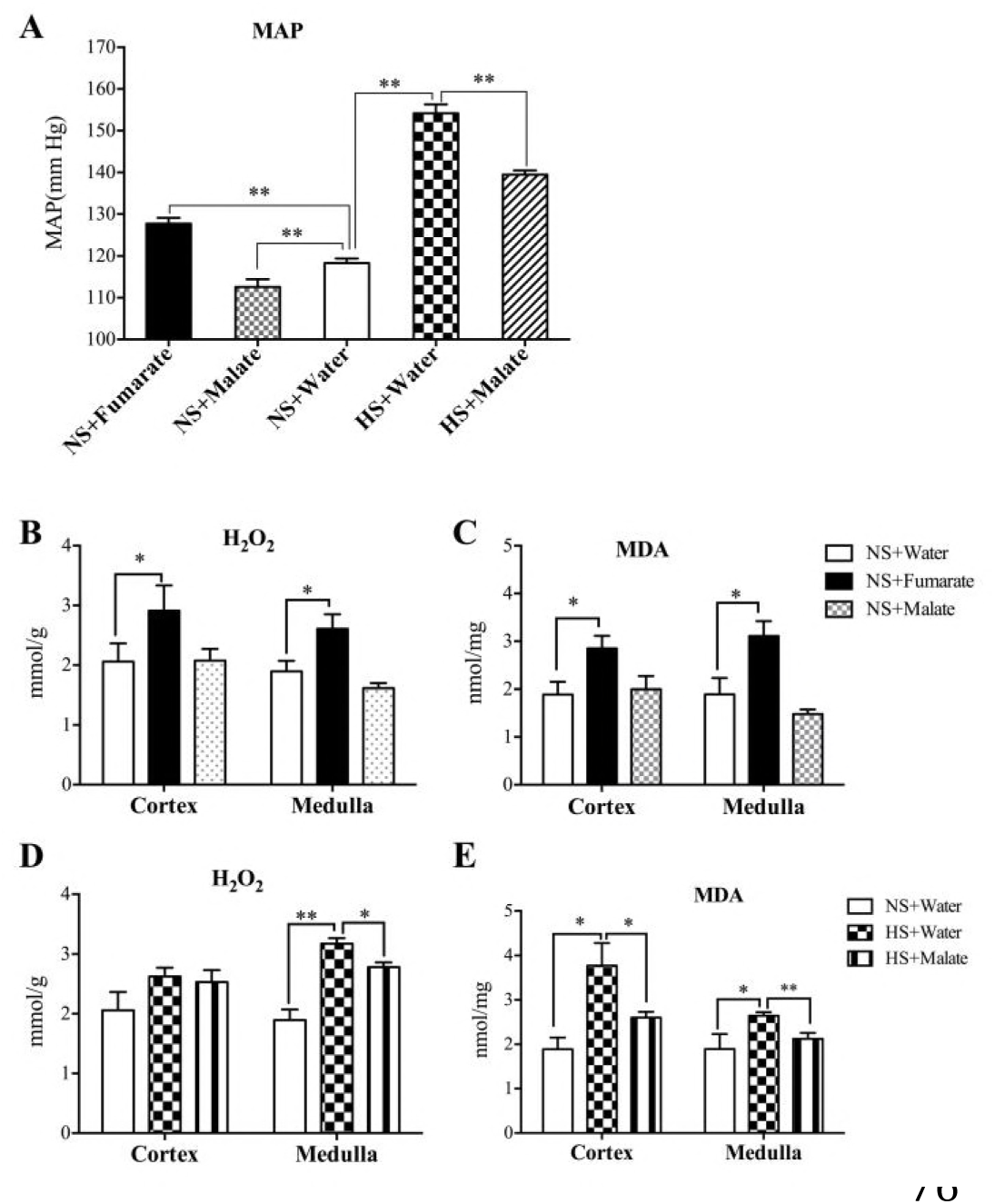
Levels of H_2_O_2_ and MAD in the kidney of SS rats with fumarate or malate supplementation. A Mean arterial blood pressure (MAP) of SS rats on 0.4% salt (normal salt, NS) diet with or without malate (fumarate) or 8% salt (high salt, HS) diet with or without malate supplementation by gavage. B Levels of H_2_O_2_ in the renal cortex and medulla of SS rats on 0.4% salt with or without malate (fumarate) supplementation by gavage. C Levels of MDA in the renal cortex and medulla of SS rats on 0.4% salt with or without malate (fumarate) supplementation by gavage. D Levels of H2O2 in the renal cortex and medulla of SS rats on 0.4% salt or 8% salt diet with or without malate supplementation by gavage. E Levels of MDA in the renal cortex and medulla of SS rats on 0.4% salt or 8% salt diet with or without malate supplementation by gavage. Data are presented as mean values ± SEM. n = 5-6, ^∗^p<0.05, ^∗∗^p < 0.01.

## Discussion

### Imbalance of redox state in FH knockdown HK-2 cells

GST catalyzes the conjugation of the reduced form of GSH to xenobiotic substrates for the purpose of detoxification. The upregulated expression and improved activity of GPx and GST suggested excessive consumption of GSH after FH knockdown.

GR catalyzes the reduction of GSSG to the sulfhydryl form GSH via the conversion of NADPH to NADP^+^, which is a critical molecule in resisting oxidative stress and maintaining the reducing environment of the cell. The main source of NADPH in the cell is from the pentose phosphate pathway (PPP) in which G6PD and 6PGD were defined as the rate-limiting enzyme. The activity of GR, G6PD and 6PGD was increased significantly as a response to FH knockdown in HK-2 cells (Figure 1). It was also consistent with the increased protein expression. Except PPP, NADPH also can be produced from other oxidoreductases, such as IDH2, which oxidizes a substrate by reducing an electron acceptor usually as NAD^+^ or NADP^+^ and producing NADH or NADPH. Indeed, the activity of NADP^+^-IDH2 increased significantly (Figure 1C). Meanwhile, the expression of other dehydrogenase such as LDHB, AKR1A1, ADH5, ALDH7A1 also increased significantly in Fh knockdown cells. These data suggested increasing the expression of dehydrogenase could provide more NADPH or NADH for the consumption of GR. In addition, oxidation of NADPH through mitochondrial electron transfer chains was decreased, owing to adrenodoxin oxidoreductase (FDXR), serving as the first electron transfer protein in all the mitochondrial P450 systems, reduced significantly. The ratio of NADPH/NADP^+^ slightly increased (Figure 2B).

As all key enzymes (GR, GS, GST, GPx, G6PD and 6PGD) involved in glutathione metabolism increased their activities or protein expression, suggested that the oxidation-reduction reaction that GSH participated in was extremely active, excessive ROS and reducing molecular produced in HK-2 cells at the same time after FH knockdown.

The ratio of GSH/GSSG is often used as an indicator of the cellular redox state since it is the most abundant thiol-disulfide redox buffer in the cell [13]. Normally, GSH is about 30-100 fold excess over GSSG. The oxidation of only a small amount of GSH to GSSG can significantly change the ratio and the redox status of the cell. However, knockdown of FH expression resulted in a significant decreasing of GSH/GSSG in HK-2 cells (Figure 2E). The results further confirmed that the insufficient FH activity could result in the oxidative stress. Nevertheless, where did these ROS come from?

The activity of NADPH oxidases was detected and showed significantly increased in FH knockdown HK-2 cells, which should be the contributor of ROS production (Figure 1A). Mitochondrial enzymes are so far known to generate ROS including the TCA cycle enzymes aconitase (ACO), OGDH, PDH, GPDH (glycerol-3-phosphate dehydrogenase), B5R and Complexes I, II and III [14]. Current differential proteomics analysis data showed that the expression of NDUFS5, an important member of the Complex I, reduced significantly. Its content of expression modulates the efficiency of mitochondrial electron transport chain and ATP level [15]. The fully assembling of Complex I will be disturbed by insufficient expression of NDUFS5, the ROS generation will be promoted [16, 17]. RISP as one of the catalytic subunits of Complex III, it had been demonstrated that deficiency of RISP influenced the stability of Complex I, Complex III and increased generation of ROS [18]. Both RISP and Cyt b-c1 complex subunit 6 reduced 1.5 fold in FH knockdown HK-2 cells. It suggested complex III contributed to the production of ROS. Meanwhile, the upregulation of both OGDH and B5R also promoted ROS production. ATP synthase had been defined as one important site of ROS generation. Its activity can modulate ROS formation and ATP level in mitochondria [19]. ATP5A1 was identified as a target for calpain-1 in diabetic hearts, leading to its proteolytic degradation, disruption of ATP5A1 may compromise mitochondrial function, resulting in excessive ROS generation in mice [20]. Current data showed that the expression of ATP synthase subunit ATP5A1, ATP5C1, ATP5I and ATP5L reduced significantly, and accordingly ATP level reduced too (Figure 1B).

Briefly, the signature of redox metabolism for FH knockdown HK-2 cells would be NOX, α-OGDH, B5R, ATPase and complex I, III worked as the main sites of ROS formation, and upregulation expression of GR, GS, GST, GPx, G6PD, 6PGD, SOD2, IDH2 and NNTH worked as an antagonistic response to the excessive ROS. The final substantial drop in GSH/GSSG ratio was the result of FH knockdown. The redox metabolism for FH knockdown HK-2 cells was summarized in Figure EV5.

### The oxidative stress in renal medulla and cortex of SS rats

It had been demonstrated that NOX was the major source of excess superoxide produced in renal medulla of SS rats [5]. The increase of NOX activity is consistent with it in FH knockout HK-2 cells. The expression of OGDH increased in renal medulla, on the contrary, the expression of ATP5A1 reduced significantly in renal medulla and cortex of SS rats compared with SS.13^BN^. These results suggested that OGDH and ATPase may also contribute to ROS formation in renal medulla of Dahl SS rats. Meanwhile, one mitochondrial proteomics analysis also suggested that Complex I and Complex III were also the ROS production sites in renal medullary thick ascending limb (mTAL) of the SS rats [12]. Briefly, B5R, NOX, α-OGDH, ATPase, Complex I and Complex III would be the sites of ROS production in renal medulla of SS rats.

For the antioxidant response of SS rats, the synthesis of GSH was also modulated by the insufficient substrates, as the levels of glycine, glutamate and cysteine were reduced in renal medulla compared with SS.13^BN^. The levels of GSH, GSSG and total GSH (GSH+GSSG) in renal medulla of SS rats were the similar to those in FH knockdown HK-2 cells. Especially, the ratio of GSH/GSSG also reduced significantly. GPx had been reported increase in SS medulla [21]. Other dehydrogenases such as IDH2, 6GPD and G6PD also increased their expression or activity. These enzymes in renal medulla of SS rats had the same response to oxidant stress as FH knockout HK-2 cells. The full comparison involved in redox balance between SS rats and FH knockout HK-2 cells was listed in Table EV5. All the data suggested that the imbalance of redox metabolism in renal medulla of SS rats was highly similar to FH knockout HK-2 cells.

### Fumarate and malate contribute to the imbalance of redox and hypertension in SS rats

There is no evidence to confirm that FH is the site of ROS production, but the excessive fumarate formed by the insufficient FH activity could lead to oxidative stress by succination of GSH [22]. This covalent adduct between fumarate and GSH reduced the efficiency and level of GSH in the cell. Meanwhile, the total GSH (GSH+GSSG), GSH reduced significantly in SS renal cells and HK-2 cells as inadequate synthetic substrate, glycine and glutamate. Obviously, the insufficient GSH flux attenuated the antioxidant ability of GSH system and promote ROS formation. Indeed, the content of fumarate had been proved to increase in SS renal medulla,

Current data had proved extra fumarate oral administration significantly increased the levels of H_2_O_2_ and MDA, and exacerbated the elevation of MAP (Figure 5). As the production of FH, excess malate oral administration could reduce the generation of ROS and attenuated the high salt diet induced hypertension. The mechanism of malate working is mainly through increasing synthesis of arginine and NO [11].

## Conclusion

The insufficient FH activity led to the accumulation of fumarate in renal medulla of SS rats, and excessive ROS generated from NOX, OGDH, ATPase and mitochondrial respiratory chain complex I, III. The increased G6PD, 6PGD, IDH and other dehydrogenase activity became the main antagonistic response of cells to oxidative stress induced by FH insufficiency. Glutathione metabolism, as the most important antioxidant mechanism, lost its balance due to insufficient synthesis substrate and resulted in the decreasing of antioxidant capacity, which made the cells under the oxidative stress state. Therefore, the defect of FH gene located on chromosome 13 of SS rats cause a wide range of changes at several sites of ROS production and antioxidant mechanisms

## Materials and Methods

### Cell culture and siRNA-mediated RNA interference

HK-2, a human kidney epithelial cell line was obtained from and cultured as suggested by Conservation Genetics CAS Kunming Cell Bank (China). RNA interference was performed as described previously [9, 23]. The small interfering RNA (siRNA) for human fumarase targets the CCAGGAUUAUGGUCUUGAUTT sequence (Gene Phaima Technologies, China). Western blot and enzyme activity analyses were performed 64 hours after transfection.

### Western blotting

Western blot was performed as described previously [24]. Sources and dilutions of primary antibodies used were Fumarase, FH (CST, USA) 1:1000; Superoxide dismutase [Mn], SOD2 (Wanleibio, China) 1:2000; NADP^+^-dependent isocitrate dehydrogenase cytoplasmic, NADP^+^-IDH1 (Wanleibio, China) 1:2000; NADP^+^-dependent isocitrate dehydrogenase, mitochondrial, NADP^+^-IDH2 (BOSTER, China) 1:2000; Heat shock protein HSP 90, HSP90 (Wanleibio, China) 1:2000; 2-Oxoglutarate dehydrogenase, OGDH (CST, USA) 1:1000; ATP synthase subunit alpha, ATP5A1 (Proteintech, USA) 1:2000; β-Actin (CW0096, China) 1:10000. Secondary antibodies were HRP-conjugated goat anti-mouse IgG (BOSTER, China) or HRP-conjugated goat anti-rabbit IgG antibody (Santa Cruz, USA). Signals were developed with Thermo Super Signal West Pico Trial Kit. The intensities of the target proteins were quantified using Image J software and β-actin was used as loading control.

### Measurement of ROS and H_2_O_2_

ROS level was detected using dichlorodihydrofluorescein diacetate (H_2_DCF-DA) (Sigma-Aldrich) [25]. Fluorescence was determined with a fluorescence spectrometer (TECAN Infinite M200 PRO, Schweiz) at 485 nm (excitation) and 538 nm (emission). Cellular oxidant levels were expressed as relative DCF fluorescence per sample protein amounts (measured using a BCA assay). H_2_O_2_ generation was detected using Hydrogen Peroxide assay kit (Jiancheng Biochemical).

### Proteomic analysis

#### (1) Protein reduce, cysteine block and digest

HK-2 cells were lysed in a buffer containing 9 mol/L Urea, 4% CHAPS, 1% DTT, 1% IPG buffer (GE Healthcare) at 4 °C for 1 h. The cell lysates were centrifuged at 15000 g for 25 min. The supernatant were collected and the protein concentration was determined by Bradford method [26]. A volume corresponding to 100 μg of protein were precipitated with 5 volumes of acetone at–20°C for 1h. After centrifugation for 10 min at 15000 g, the deposits were collected and dried by vacuum freezing dryer. Then dissolve protein pellets in 50 μL iTRAQ dissolution buffer (Applied Biosystems) and add 4 μL Reducing Reagent (Applied Biosystems) at 60 °C for 1 h. After cooling samples to room temperature (RT), cysteine residues were blocked with 2 μL 200 mmol/L methylmethanethiosulfate by incubating at RT for 10 min. The protein solutions were added to 10 kDa ultrafiltration tubes and cleaned by centrifugation (12000 rpm, 20 min). Then 100 μL iTRAQ dissolution buffer was added in ultrafiltration tubes, centrifuged at 12000 g for 15 min and repeat this step three times. Place the column in a new tube, add 50 μL sequencing-grade trypsin (50 ng/μL) and incubate at 37 °C for 12 hours. Afterward, centrifuge by 12000 g for 20 min and collect the peptide. Transfer the filter units to new collection tube and add 50 μL iTRAQ dissolution buffers to centrifuge the tube again. Combined the two filter solutions.

#### (2) Protein digestion and labeling with iTRAQ reagents

The labeling reactions were performed following the manufacturer’s recommendations. Peptide samples from negative control HK-2 cells were labeled with iTRAQ reagent, 113 isobaric tag and peptide samples from FH siRNA-treated HK-2 cells were labeled by adding the same amount of iTRAQ reagent, 114 isobaric tag. After 2 h of incubation at RT, reactions were stopped by adding 100 μL of water, to each vial.

#### (3) Peptide fractionation by strong cation exchange chromatography

The mixed peptides were fractionated by strong cation exchange chromatography (SCX). Samples were separated using an Agilent 1200 HPLC System (Agilent), Michrom column (Poly-SEA 5μ 300Å 2.0 × 150 mm) at a flow rate of 0.3 ml/min, using a nonlinear binary gradient starting with buffer A (10 mmol/L formic acid, 20% acetonitrile) and transitioning to buffer B (500 mmol/L formic acid, 20% acetonitrile). First, the column was washed with buffer A for 5 min and peptides were eluted with a four-step gradient: first a linear gradient of 5-50% buffer B for 25 min, followed by a linear gradient of 50-80% buffer B for 5 min. The gradient was ramped to 100% buffer B in 1 min, and held for 10 min. Collect the first segment from 0-5 min, then collect each segment with 4 min interval for the 6-44 min, and for the last segment from 45-50 min, with a total of 12 segments. Dry every segment in a vacuum freezing dryer for LC-MSMS analysis.

#### (4) RPLC-MS/MS analysis

The online Nano-RPLC was employed on the Eksigent nanoLC-Ultra^™^ 2D System (AB SCIEX). The samples were loaded on C_18_ nano LC trap column (100 μm×3 cm, C_18_, 3 μm, 150 Å) and washed by Nano-RPLC Buffer A (0.1% formic acid, 2% acetonitrile) at 2μL/min for 10 min. An elution gradient of 5-35% acetonitrile (0.1% formic acid) in 70 min gradient was used on an analytical Chrom XP C_18_ column (75 μm×15 cm, C_18_, 3 μm 120 Å) with spray tip. Data acquisition was performed with a Triple TOF 5600 System (AB SCIEX, USA) fitted with a Nanospray III source (AB SCIEX, USA) and a pulled quartz tip as the emitter (New Objectives, USA). Data were acquired using an ion spray voltage of 2.5 kV, curtain gas of 30 PSI, nebulizer gas of 5 PSI, and an interface heater temperature of 150 °C. For information-dependent acquistion (IDA), survey scans were acquired in 250 ms and as many as 35 product ion scans were collected if they exceeded a threshold of 150 counts per second (counts/s) with a 2^+^ to 5^+^ charge-state. The total cycle time was fixed to 2.5 s. A rolling collision energy setting was applied to all precursor ions for collision-induced dissociation (CID). Dynamic exclusion was set for ½ of peak width (18 s), and the precursor was then refreshed off the exclusion list.

#### (5) Protein identification and quantification

Data were processed with Protein Pilot Software v. 4.0 (AB SCIEX, USA) against Homo sapiens database using the Paragon algorithm [27]. Protein identification was performed with the search option: emphasis on biological modifications. The database search parameters were as following: instrument was Triple TOF 5600, iTRAQ 2-plex quantification, cysteine modified with iodoacetamide, biological modifications were selected as the ID focus, trypsin digestion. An automatic decoy database search strategy was employed to estimate the false discovery rate (FDR) using the PSPEP (Proteomics System Performance Evaluation Pipeline Software, integrated in the Protein Pilot Software). The FDR was calculated as the false positive matches divided by the total matches. The iTRAQ 2-plex was chosen for protein quantification with unique peptides during the search. A total of 3174 proteins were identified in HK-2 cells, with the value of global FDR from fit less than 1% and identified with more than two peptides for 95% confidence were considered for further analysis. Proteins with a fold change of >1.50 or <0.67 were considered to be significantly differentially expressed.

### GO and KEGG pathway enrichment analysis

The multi-omics data analysis tool, Omics Bean, was used to analyze the obtained proteomics data (http://www.omicsbean.com:88), in which distributions in biological functions, subcellular locations and molecular functions were assigned to each protein based on Gene Ontology (GO) categories. The Kyoto Encyclopedia of Genes and Genomes (KEGG) pathway analysis was performed to enrich high-level functions in the defined biological systems.

### Enzymes activities assays

The obtained pellets were lysed by three freeze–thaw cycles to ensure that the mitochondrial membrane was disrupted and enzymes were accessible. The protein content was determined with BCA Protein Assay Kit (Beyotime Institute of Biotechnology, China).

The enzyme activity of FH was measured based on a previous reported method [8]. The activities of glutathione reducatase (GR) were assayed using the Beyotime Detection kits (Beyotime Institute of Biotechnology, China) according to the corresponding protocols. The activities of glutathione S-transferase (GST), glutathione peroxidase (GPx) in HK-2 cells were assayed using the Jian cheng Biochemical detection kits according to the standard protocols.

Glucose-6-phosphate dehydrogenase (G6PD) and 6-phosphogluconate dehydrogenase (6PGD) activities assay were performed as described previously [28, 29]. Isocitrate dehydrogenase, cytoplasmic (NADP^+^-IDH1) and isocitrate dehydrogenase, mitochondrial (NADP^+^-IDH2) are NADP^+^-dependent isocitrate dehydrogenase (NADP^+^-IDH). NADP^+^-IDH activity was assayed by spectrophotometric monitoring of the reduction of NADP^+^ to NADPH at 340 nm according to protocol by Connie S. Yarian [30]. The NADPH oxidase activity was determined using the NADPH oxidase assay kit purchased from Nanjing Jiangcheng Bioengineering Institute according to the manufacturer’s instructions.

#### GSH (GSSG) and NADPH (NADP^+^) measurements

The determination of reduced glutathione (GSH) and oxidized glutathione (GSSG) were performed using DTNB method with kits according to the corresponding kit protocols (Beyotime, China). The determination of NADP^+^ and NADPH were performed with kits according to the corresponding kit protocols (Comin, China).

#### GC-MS assay

The extraction of metabolites in tissue and cells samples was performed according to the method described in previous reports [31, 32]. Dried tissue and cell residues were derivatized using a two-step procedure with a minor modification [33, 34]. For oximation, 80 μL of 10 mg/mL methoxyamine hydrochloride (Sigma-Aldrich) dissolved in pyridine was mixed with a lyophilized sample, and kept at 30°C for 90 min. Then, 80 μL of N,O-bis(trimethylsilyl)-trifluoroacetamide (BSTFA) with 1% TMCS (Sigma-Aldrich) was added for derivatization, and heated to 70°C for 60 minutes. GC-MS analysis was performed according to our previous report [33], using a 7890A GC/5975C Inert MSD (Agilent Technologies, Wilmington, DE), coupled with a DB-5 column (30 m × 0.25 mm I.D.; film thickness: 0.25 μm; Agilent J&W Scientific, USA). Helium was used as carrier gas at a constant flow rate of 1 ml/min. The GC temperature programming was set to 2 min isothermal heating at 80°C, followed by 10°C/min oven temperature ramps to 120°C, 5°C/min to 260°C, and then increased at a rate of 10 °C /min to 300°C, where it was held for 2 min. Fumarate, L-malate, glutamate, glycine, L-cysteine standards were analyzed under identical experimental conditions for further identification. One μL of the final derivatized aliquots was injected into the GC-MS.

#### Measurement of ATP

The ATP assay was performed with a kit from Promega (G7570) according to manufacturer’s instruction. The luminescence was recorded in a TECAN Infinite M200 PRO (Schweiz) with an integration time of 5s per well.

#### Animals and tissues

Male SS rats (SS/JrHsdMcwi) and consomic, salt-insensitive SS.13^BN^ rats [35] were bred in a pathogen-free animal house and maintained on a purified AIN-76A rodent diet (Dyets) containing 0.4% NaCl with free access to water. For tissue collection, kidneys were flushed in situ with cold saline and the renal cortex and medulla were quickly removed and snap-frozen in liquid nitrogen and stored at −80°C. Frozen tissues were used for preparation of tissue homogenates to perform activity assays, Western blots and metabolite analysis. The experiments were approved by the Institutional Animal Ethics Committee of Xi’an Jiaotong University.

#### Fumarate and malate supplementation

Male SS rats were fed a 0.4% NaCl diet since weaning. After three days of stable baseline blood pressure at approximately 7 weeks of age, rats were switched to an 8% NaCl diet (HS). Fumarate (50 mg/kg/d) [36], malate (600 mg/kg/d) [37], or an equal volume of distilled water was delivered by gavage. Serum and kidneys were harvested at the end of blood pressure measurement on day 17 of the HS diet.

#### Measurement of blood pressure

Blood pressure of conscious rats was measured using tail-cuff plethysmography with a CODA-4 computerized system (Kent Scientific Corporation, Torrington, Connecticut, USA). Mean arterial pressure (MAP) was measured between 14:00 and 16:00 by a single experienced operator. Rats were trained for the procedure for five consecutive days. After calibrating the tail-cuff apparatus, rats were placed on the platform and allowed to rest in a glass restrainer. A black conical plastic piece with a nose opening was placed over the head region of the rat to cover the eyes and allow the rat to rest. The platform was heated to 30–32 °C to increase the detection of the oscillation waveforms generated by the blood flow in the tail. Each session was composed of 15 successive measurements of the systolic and diastolic pressure, which were averaged. Heart rate was recorded at the same time.

#### Measurement of H_2_O_2_ and MDA in the kidney of SS rats

H_2_O_2_ and malondialdehyde (MDA) generation were detected using Hydrogen
Peroxide assay kit (Jiancheng Biochemical).

#### Statistical analysis

All values are presented as mean ± SEM. All data were analyzed using the Student’s t-test or ANOVA analysis. Probabilities of < 0.05 were considered to be statistically significant. All of the statistical tests were performed with the Graph Pad Prism software, version 6.0 (Graph Pad Software Inc., San Diego, CA).

## Acknowledgments

This study was supported in part by the National Natural Science Foundation of China (Grant No. 81570655, 81770728, 51703178)

## Expanded View Figure legends

**Figure EV1 - Down-regulation of fumarase expression and activity is associated with the accumulation of fumarate, ROS and H_2_O_2_ in the HK-2 cells.**

A Western bloting densitometric analysis of fumarase of HK-2 cells on both negative control (NC) and FH-siRNA treated (FH-SiRNA) with the associated gel shown.

B Fumarase (FH) activity.

C Fold changes of fumarate and L-malate level in fumarase insufficient HK-2 cells were calculated as a ratio relative to negative control.

D ROS levels.

E, H_2_O_2_ levels. Mock siRNA targeting fumarase that had no sequence homology with any mammalian genes as a negative control.

Data are presented as mean values ± SEM, n = 4-6, ^∗^p < 0.05, ^∗∗^p < 0.01 compared to negative control.

**Figure EV2** - Expressional changes of SOD2, IDH1, IDH2 and HSP90 in the HK-2 cells induced by fumarase insufficiency.

Left: a representative result of western blotting shows the expressions of SOD2, IDH1, IDH2, and HSP90 in fumarase insufficient HK-2 cells compared to negative control.

Right: histogram shows the expression levels of the four proteins in these HK-2 cells as determined by densitometric analysis.

β-actin is used as the internal loading control. Mock siRNA targeting fumarase that had no sequence homology with any mammalian genes as a negative control.

Data are presented as mean values ± SEM. n=4, ^∗^p < 0.05, ^∗∗^p < 0.01 significant difference from negative control.

**Figure EV3 - GO annotation of identified significant changes proteins in three categories: biological process (BP), cellular component (CC) and molecular function (MF).**

All of biological processes were ranked in term of the enrichment of the differentially expressed proteins, and the top 10 are presented.

**Figure EV4 - Distribution of enriched KEGG Pathway. Top 10 enriched pathway are shown here.**

P value =0.01 (red) and P value =0.05 (blue) as two selected cutoff are highlighted on the figure, as an indicator to show how significant the results are based on KEGG enrichment.

**Figure EV5. Role of fumarase insufficiency in reactive oxygen species metabolism.**

The proteins or compounds involved in ROS generation (red stars) and antioxidant defence (blue stars). The red arrows indicate proteins or compounds identified as significantly increased in our study. The green arrows indicate proteins or compounds identified as significantly decreased in our study. FH, Fumarase; OGDH, 2-oxoglutarate dehydrogenase; C I, C II and C III, the electron-transport chain (ETC) complexes I, II and III; B5R, cytochrome b5 reductase; NNTH, Nicotinamide nucleotide transhydrogenase; G6PD, Glucose-6-phosphate 1-dehydrogenase; 6PGD, 6-phosphogluconate dehydrogenase; IDH2, Isocitrate dehydrogenase [NADP+], mitochondrial; PRDX5, Peroxiredoxin-5, mitochondrial; FDXR, NADPH:adrenodoxin oxidoreductase, mitochondrial; SOD2, Superoxide dismutase [Mn]; GS, Glutathione synthetase; GR, Glutathione reductase; GPx, Glutathione peroxidase; GSH, reduced glutathione; GSSG, oxidized glutathione.PM, Plasma membrane; OMM, Outer mitochondrial membrane; IMM, Inner mitochondrial membrane.

## Expanded View Table legends

**Table EV1.** Summary of the total proteins were identified and quantified reliably by iTRAQ.

**Table EV2.** Differentially expressed proteins in FH-siRNA treated HK-2 cell line identified from iTRAQ analysis.

**Table EV3.** GO enrichment of the total significant changes proteins.

**Table EV4.** KEGG enrichment of the total significant changes proteins.

**Table EV5.** The enzymes and metabolites involved in ROS generation and antioxidant defence in fumarase insufficient HK-2 cell and SS rats.

